# Human Liver Organoids as a Patient-derived Model for HBV Infection and Cellular Response

**DOI:** 10.1101/2022.10.20.513112

**Authors:** Chuan Kok Lim, Ornella Romeo, Andrew P Chilver, Bang Manh Tran, Dustin J Flanagan, Emily N Kirby, James Breen, Elizabeth Vincan, Nadia Warner, Erin M McCartney, Mark B Van Der Hoek, Andrew Ruszkiewicz, Edmund Tse, Michael R Beard

**Affiliations:** Research Centre for Infectious Diseases, School of Biological Sciences, The University of Adelaide, SA, Australia; Royal Adelaide Hospital, Adelaide, Australia; SA Pathology, Adelaide, Australia; Department of Infectious Diseases, The University of Melbourne at the Doherty Institute for Infection and Immunity, Melbourne, VIC 3000, Australia; Victorian infectious Diseases Reference Laboratory, Royal Melbourne Hospital at the Doherty Institute for Infection and Immunity, Melbourne VIC 3000, Australia; Monash Biomedicine Discovery Institute, Monash University, Clayton, Australia; South Australian Health and Medical Research Institute (SAHMRI), Adelaide, Australia; Curtin Medical School, Curtin University, Perth, WA 6102, Australia; Gastroenterology and Hepatology, Royal Adelaide Hospital, Adelaide, Australia

**Author notes:** Correspondence to be addressed to, Michael R Beard PhD, Address: Viral Pathogenesis Research Laboratory, Molecular Life Sciences, University of Adelaide, North Terrace, Adelaide, Australia, Chuan Kok LIM MBBS PhD, Current Address: Victorian Infectious Diseases Reference Laboratory and the University of Melbourne at the Doherty Institute of Infection and Immunity, Melbourne, VIC 3000, Australia. **Authors’ Contributions**, C.K.L, N.W, S.L and M.R.B performed conceptualization and supervision of the study. C.K.L and M.R.B arranged funding acquisition. C.K.L, O.R, A.P.C, and E.K developed methodology and investigation. C.K.L, E.M. E.T, A.R involved in project administration. B.M.T, E.V. E.M, E.T, A.R, S.L provided resources and method development. J.B, C.K.L and A.P.C performed data curation and formal analysis. C.K.L and M.R.B prepared visualization, writing original draft, review and editing.

**Keywords:** Liver organoids, LGR5, HBV, NTCP, Interferon-stimulated genes, antivirals

## Abstract

**Background & Aims:** Current HBV *in vitro* model systems suffer from many physiological limitations that restrict understanding of complex viral-host interactions and thus prohibit prediction of disease *in vivo*. We developed and assessed adult stem cell (AdSC) derived liver organoids as a novel model system for characterisation of the HBV lifecycle, the cellular response to infection and demonstrate their utility in assessing antiviral and immunomodulator response. This model system has the potential to be used in predicting individual HBV responses to antivirals and viral reactivation in the setting of immunosuppressive agents.

**Methods:** Ductal stem cells were isolated from healthy tissue acquired from liver resections or biopsy (n=12). Wnt3a & RSPO-1 containing medium was used to stimulate ductal stem cell expansion into organoids which were subsequently differentiated into hepatocyte-like cells. Mature hepatocyte metabolic markers (albumin, CYP3A4) and HBV entry receptor (Na-taurocholate co-transporting polypeptide, NTCP) expression were evaluated throughout differentiation using qRT-PCR and confocal microscopy. We assessed the organoids culture conditions required for HBV infection and HBV life cycle using HepAD38 (genotype D) and plasma derived HBV (genotype B & C). HBV infection was confirmed using immunofluorescence staining (HBcAg), qRT-PCR (RNA, cccDNA, extracellular DNA) and ELISA (HBsAg and HBeAg). We also assessed drug responsiveness using antivirals and an immunosuppressive agent, and cellular responses (interferon-stimulated genes) using interferon-α and viral mimic (PolyI:C).

**Results:** Following differentiation, organoids underwent structural remodelling and changes in cellular polarity, accompanied with an increase in albumin, CYP3A4 and NTCP mRNA expression. Optimal HBV infection was achieved in well-differentiated organoids using spinoculation of at least 200 copies/cell of AD38 derived HBV. Infected organoids demonstrate time and donor dependent increase in HBV RNA, cccDNA, extracellular DNA, HBe and HBsAg consistent with viral replication and antigen secretion. Using these markers we assessed drug-responsiveness to the HBV entry inhibitor, Myrcludex B and the JAK inhibitor, Baricitinib. Despite having a very robust interferon stimulated gene response to interferon-α and PolyI:C stimulation, HBV infection in liver organoids did not reveal innate immune activation.

**Conclusions:** AdSC derived liver organoids support the full life cycle of HBV with significant donor dependent variation in viral replication and cellular responses. These features can be utilised for development of personalised drug testing platform for antivirals.

**Lay Summary:** Human liver organoid culture provides a personalised assessment of HBV infection, replication and responsiveness to antiviral therapy. This model system has a robust innate immune response and could be used to assess novel immune-modulating curative therapy.

## Introduction

Chronic or persistent Hepatitis B virus (HBV) infection is the leading cause of liver cirrhosis and hepatocellular carcinoma (HCC) and in 2015 it was estimated to affect more than 250 million people worldwide with more than 800,000 deaths^1^. At present, there is no effective curative therapy for chronic hepatitis B, although there is a major global initiative to identify strategies for a HBV cure. However, attempts to develop curative therapy and a greater understanding of the response to infection have been hampered by a lack of satisfactory *in vitro* model systems that faithfully mimics the physiology of hepatocytes and the liver.^2^ Primary human hepatocytes (PHH) are regarded as the “gold-standard” model for *in vitro* HBV infection, however they rapidly dedifferentiate in culture, are costly and have a limited lifespan that hampers their use in modelling for chronic HBV infection.^3^ Ectopic expression of NTCP, the HBV cellular receptor in tumour derived cell lines such as HepG2 cells has been used extensively in studying HBV infection *in vitro* but suffers from altered physiology and cellular response.^4^ Importantly, this approach is non-physiological and represents an over-simplification of HBV infection. Furthermore, it overlooks genetic variation of hepatocytes, thus limiting its utility in predicting clinical response and the development of a personalised medicine approach to HBV infection management.

Human liver organoids developed by *Huch et al*. consist of self-organising hepatic LGR5+ stem cells which can be expanded long-term with genetic stability.^5^ They exhibit mature hepatocyte and ductal phenotypes following differentiation. Most importantly, they retain host genetics for the liver from which they were derived that allows for personalised drug testing and investigation of variance in host response to infection. Unlike many other stem-cell or primary hepatocyte derived model systems, the LGR5+ human liver organoid culture does not involve the process of parental cell reprogramming to achieve multi-potency for long-term expansion, thus limiting the introduction of mutations in culture. ^5^ This quality is particularly important for potential clinical applications in the area of autologous transplantation. In addition, the LGR5+ liver organoid culture is the first *in vitro* culture system that exhibits self-organising capability of its cellular structures to mimic the host organ, sometimes referred to as four-dimensional (4D) culture. This contrasts with traditional culture model systems using cell monolayers (including primary hepatocytes) or 3D culture which requires a structural framework to support cell growth, with both representing significant departure from *in vivo* conditions.

In this study, we developed human liver organoid cultures derived from either core liver biopsies or liver resections as a physiological model for HBV infection. We assessed the characteristics of human liver organoids that are important for HBV infections, in particular Na-taurocholate co-transporting polypeptide (NTCP) and other HBV-dependent host-factors.^6^ We also examined the relevance of structural changes and polarity in human liver organoids, which were previously shown to be important for productive HBV trafficking and export.^7^ Human liver organoids expressed NTCP, were permissive to HBV infection and are innate immune competent that provides a primary model system to assess HBV antiviral responses and possible genetic variation to HBV infection and the cellular response.

## Materials and Methods

### Patient Selection and Liver Organoids Isolation

Patients who underwent liver resections for metastatic carcinoma or core biopsies were recruited through the Gastroenterology and Liver Tissue Repository at the Royal Adelaide Hospital with ethics approval from the hospital research committee. Tissue distant from resected tumour (>10cm away) or fragments of core biopsied tissues (weight 20-50mg) were used for organoids isolation. Tissues were minced and digested with collagenase to single cells before embedded in reduced growth factor extracellular matrix (Cultrex BME2, R&D Systems). Selective isolation of ductal stem cells was achieved using Isolation Medium (IM) consisting of Expansion Medium (EM) supplemented with 5%v/v Noggin conditioned medium (homemade), 30% v/v Wnt3a conditioned medium and 10μM of Y-27632 (Sigma) until organoids were visible. Organoids were subsequently cultured and passaged in EM which consists of Advanced DMEM/F12 (ThermoFisher) supplemented with 1x B27 (ThermoFisher), 1x N2 (ThermoFisher) 1.25mM N-acetylcysteine (Sigma), 10mM nicotinamide (Sigma), 10% v/v R-spondin conditioned medium (homemade), 10nM recombinant human gastrin (Sigma), 50ng/mL EGF (Peprotech), 100ng/mL FGF-10 (Peprotech), 25ng/mL HGF (Peprotech), 10μM forskolin (Tocris), and 5μM A83-01 (Tocris). The cell lines used to generate Wnt3a, R-spondin and Noggin conditioned media were kindly provided by Professor Hans Clevers and Professor Nick Barker.^8, 9^

### Total Cellular RNA Isolation and qPCR

Organoids were washed with cold medium to remove extracellular matrix before RNA isolation using TRIsure (Bioline) according to manufacturer’s instructions. cDNA first strand synthesis and amplification were performed using Luna Universal qPCR mastermix (NEB) with previously described qPCR conditions on StepOne Plus Real-time PCR System (Applied Biosystems).^10^ Primer sequences are provided in the supplementary material (Table 1).

**Table 1.**
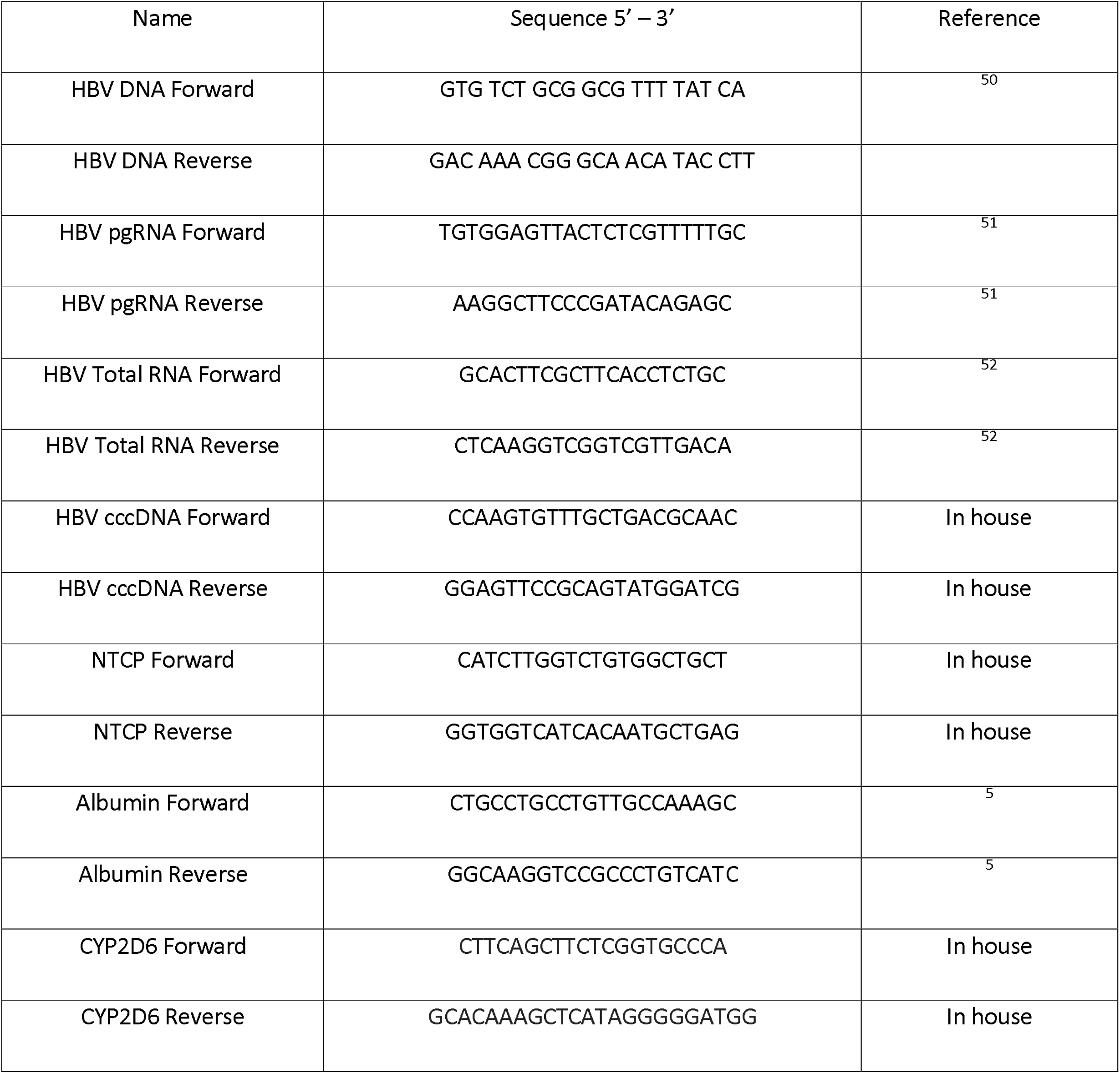
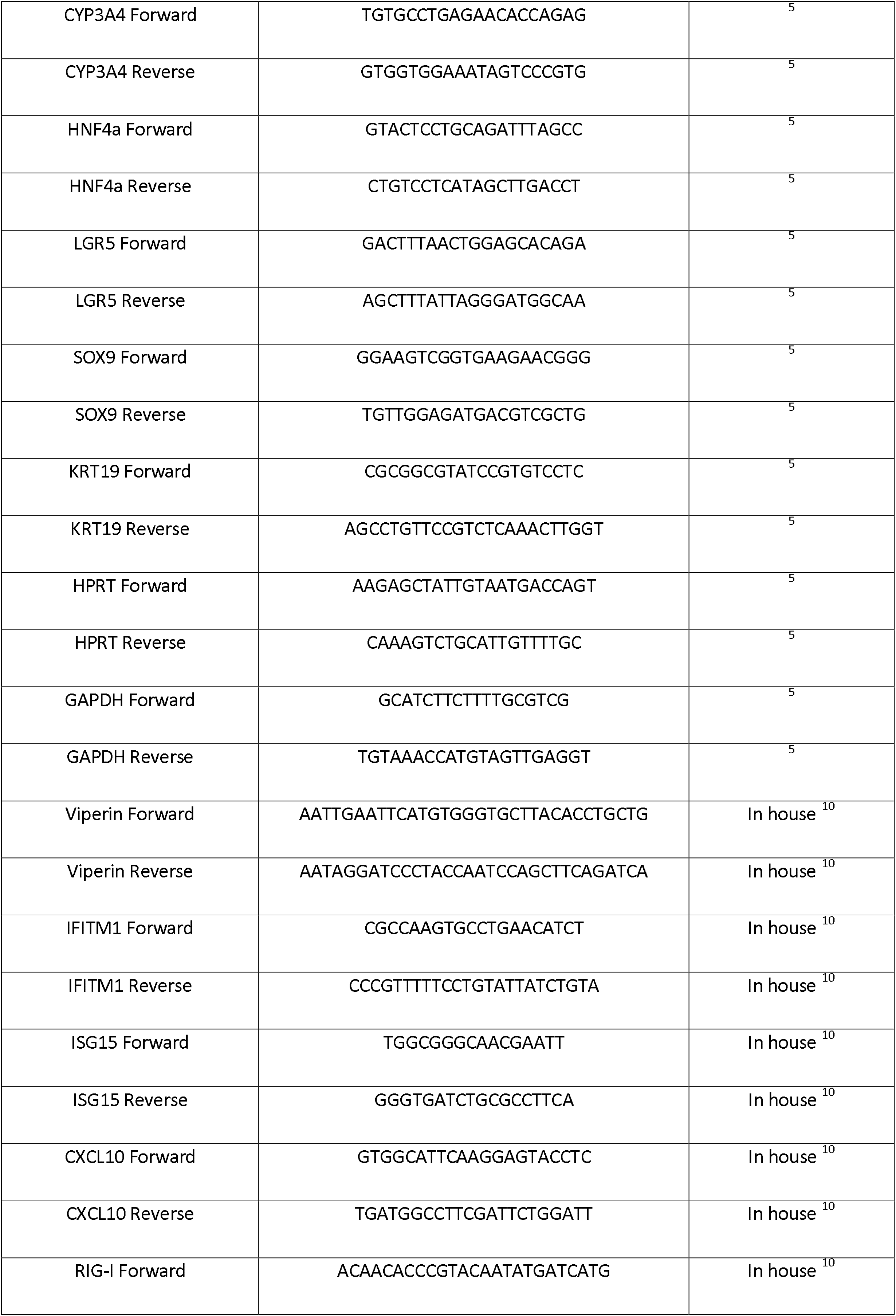
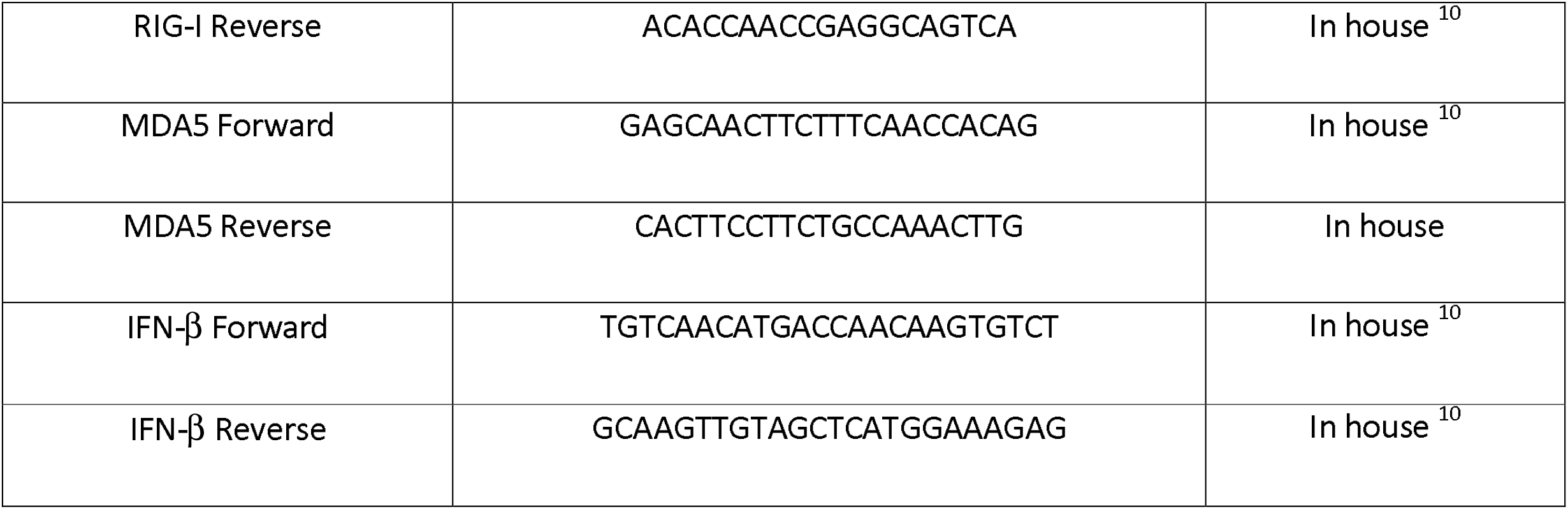
DNA Oligonucleotides

### Preparation of HBV Inoculum

Cell culture derived HBV (genotype D) was prepared from supernatant of HepAD38 cells^11^ cultured in Advanced DMEM/F12 (ThermoFisher) supplemented with 10% FCS and 400μg/mL G-418 for ten days. Culture supernatant was passed through a 0.4 μm filter and precipitated using PEG8000 (final concentration 8%) overnight before centrifuged at 15,000xg at 4°C for one hour. Viral pellet was reconstituted in Advanced DMEM/F12 at 1/50-1/100 of the original volume. Human plasma HBV was acquired from patients with high HBV viral load and stored unprecipitated in −80°C as viral stock. Genotyping was performed using S gene sequencing described by Li et al.^12^

### Preparation of Liver Organoids for HBV Infection

Following digestion with TrypLE (ThermoFisher), organoids were seeded as single cells and cultured with EM supplemented with 10μM Y-27632 for 4-5 days to avoid anoikis. Once organoids reformed (20-30μm diameter), medium was changed to EM supplemented with BMP-7 (Peprotech) for 5 days, followed by 10 days in Differentiation Medium (DM) to induce hepatocyte differentiation. DM consists of Advanced DMEM/F12, 1x B27 (ThermoFisher), 1x N2 (ThermoFisher), 1mM N-acetylcysteine (Sigma), 10nM recombinant human gastrin (Sigma), 50ng/mL EGF (Peprotech), 25ng/mL HGF (Peprotech), 100ng/mL FGF-10, 0.5uM A83-01 (Tocris), 10uM DAPT (Sigma), 3μM Dexamethasone (Sjgma), 25ng/mL BMP-7 (Peprotech), and 100ng/mL (FGF-19 (Peprotech). For HBV infection, BME2 was first removed by repeated washing with cold medium before addition of HBV inoculum and DM/4% PEG8000 at 1:1 ratio. Plate spinoculation was performed at 600xg for one hour at 35°C in a non-adherent plate, before the addition of R-Spondin (10% vol/vol) and incubation for a further 24-48 hours. Organoids were washed nine times before being re-seeded in fresh non-suspension plate and cultured with DM/Y-27632.

### Quantification of HBV RNA & DNA

Extracted cellular RNA was quantified using Nanodrop 2000 (ThermoFisher) and 50ng of RNA was used in each qPCR reaction and compared with a standard curve generated from linear regression analysis of 10-fold dilutions of a linearised 1-mer HBV plasmid. Control without reverse transcription was used to rule out DNA contamination.

Supernatant of infected organoids were subjected to two DNaseI treatment (2U/μL) with 1μL MgCl_2_ prior to extraction. Intracellular and supernatant HBV DNAs were isolated using the Nucleospin Tissue extraction kit (Macherey-Nagel) following manufacturer’s instructions. Quantification was performed as described earlier. For cccDNA qPCR, samples were first digested with T5 exonuclease (NEB) with pre- and post-treatment of rcDNA as digestion control. The qPCR conditions are: 95°C, 10 min 1 cycle, 40 cycles of 95°C 15s, 60°C 5s, 72°C 45s, 88°C 2s. The cccDNA primers were validated against previously described primers by Singh et al.^13^

### RNA Sequencing of Liver Tissue and Liver Organoids

Total RNA was converted to strand specific Illumina compatible sequencing libraries using the Nugen Universal Plus mRNA-Seq library kit from Tecan (Mannedorf, Switzerland) as per the manufacturer’s instructions (MO1442 v2). Briefly, 500ng of total RNA was polyA selected and the mRNA fragmented prior to reverse transcription and second strand cDNA synthesis using dUTP. The resultant cDNA is end repaired before the ligation of Illumina-compatible barcoded sequencing adapters. The cDNA libraries were strand selected and PCR amplified for 12 cycles prior to assessment by Agilent Tapestation for quality and Qubit fluorescence assay for amount. Sequencing pools were generated by mixing equimolar amounts of compatible sample libraries based on the Qubit measurements. Sequencing of the library pool was performed using Illumina Nextseq 500 using single read 75bp (v2.5) sequencing chemistry.

Illumina high-throughput sequencing data was processed using a standard RNA-Seq analysis workflow. Raw single-end FASTQ reads were assessed for quality using *FastQC*^14^ and *ngsReports*^15, 16^, and then aligned to the GRCh37.p13 version of the human genome^17^ using the transcriptome algorithm STAR^18^. After alignment, mapped sequence reads were summarised to GRCh37 gene annotation using *featureCounts*, available through the package *RSubread* (https://bioconductor.org/packages/release/bioc/html/Rsubread.html). Differential gene expression was carried out using R/Bioconductor packages *limma*^19^ and *edgeR*^20^, using mean-variance relationship estimates of the log-counts from the limma *voom* method^21^ and contrasts defined between each sample group. KEGG pathway and Gene Ontology enrichment were also carried out using R/Bioconductor packages, and heatmaps and upset produced using *pheatmap* (https://cran.r-project.org/web/packages/pheatmap/index.html) and *upSetR* (https://cran.r-project.org/web/packages/UpSetR/index.html). All code carried out in the study is available in the RMarkdown document.

### Immunofluorescence and Immunoblotting

Organoids cultured in μ-Slide 8 well (Ibidi) were washed repeatedly with cold PBS to remove BME2 before fixation with 4% PFA for 20 minutes, followed by a 20-minute incubation with 50mM NH_4_Cl to minimise autofluorescence, and permeabilization with 0.1% Triton-X 100 for 10 minutes. After washing with cold PBS, organoids were blocked with 5% BSA/PBS for 1 hour. Next, organoids were incubated with primary antibodies diluted in 1% BSA, at 4°C for 16-48 hours on a gentle shaker, followed by incubation with secondary antibodies at 1:200 dilutions for a further 2-4 hours. Nuclear staining was performed by incubating with DAPI at 1:1000 dilutions for 15 minutes. Images were taken using the Olympus FV3000 confocal laser scanning microscope and analysed with CellSens. 3D reconstruction and deconvolution were performed using Imaris software (Bitplane). Immunoblotting was performed as previously described.^22^ Primary and secondary antibodies used are listed in supplementary materials (Table 2).

**Table 2.**
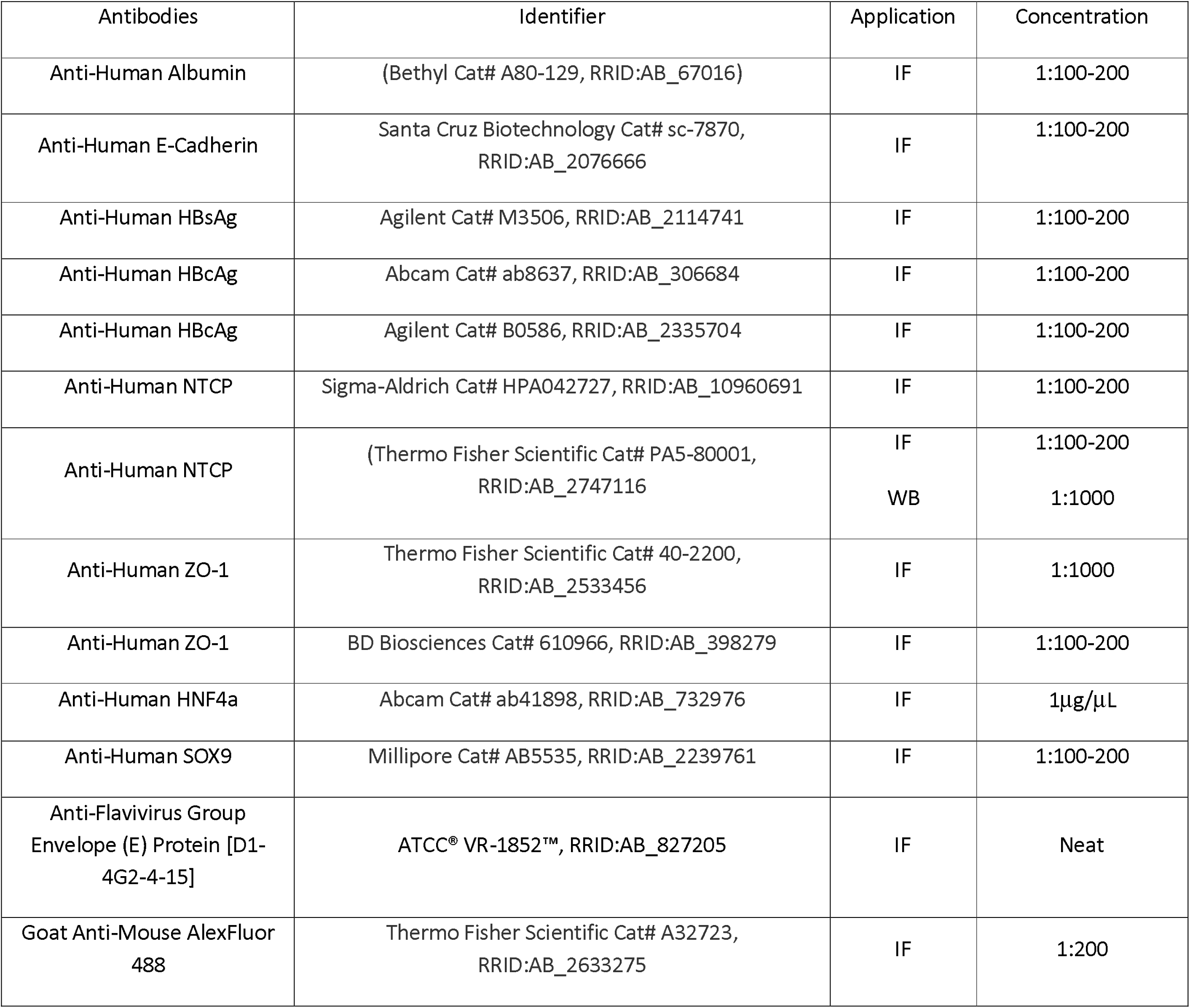
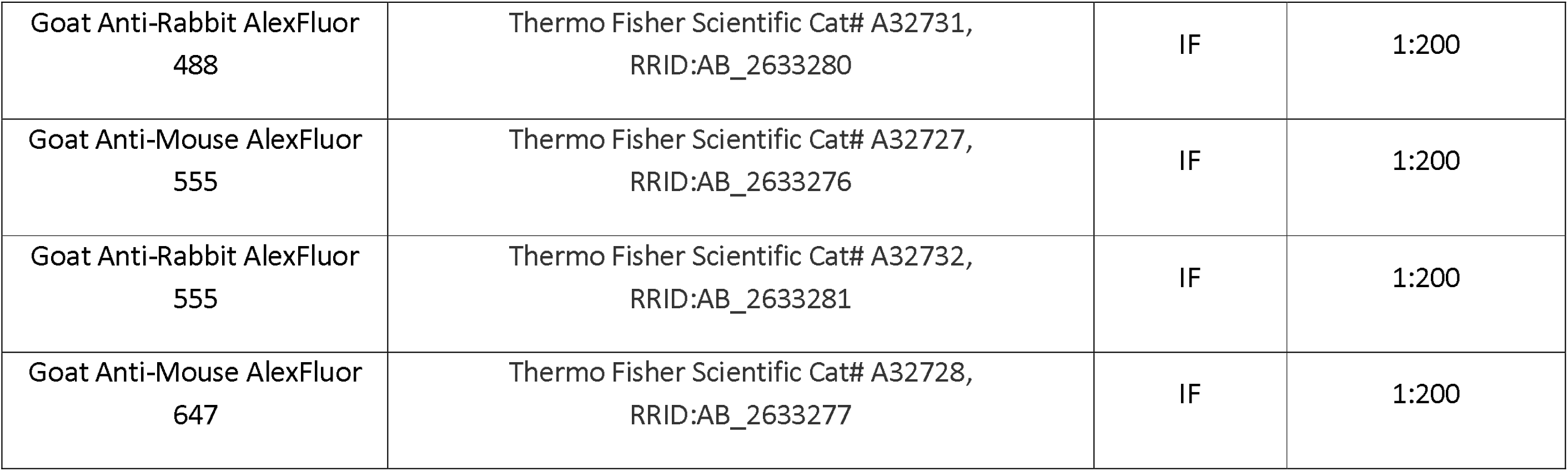
Antibodies

### Detection of secreted HBeAg and HBsAg

Supernatant (200μL) was collected from infected organoids for HBeAg and HBsAg quantification using the Elecsys HBeAg and HBsAg II kit chemiluminescent microparticle immunoassay (CMIA) on the Cobas e602 instrument (Roche).

### Transmission Electron microscopy (TEM)

Organoids were washed with cold medium to remove BME2 and fixed with 4%PFA/1.25% glutaraldehyde/4% sucrose for 1 hour. After fixation, organoids were treated with 2% OsO4 for 1 hour, washed with 4% sucrose, followed by gradual dehydration with 70, 90 and 100% ETOH and gradual infiltration from 100% propylene oxide to 100% resin before polymerization at 70°C for 24-48 hours. Blocks were sectioned at 40nm slices and imaged using the FEI Tecnai G2 Spirit TEM.

### Statistical Analysis

For qRT-PCR analysis, differential expression of genes between 2 or more groups were compared using 2-way ANOVA analysis in Prism 8. Charts were presented as mean +/− SEM with level of significance according to standard GraphPad format for p values.

### Ethical Statement

Written consent has been obtained from all patients involved in this study. The study protocol conforms to the ethical guidelines of the 1975 Declaration of Helsinki as reflected in a priori approval by the Royal Adelaide Hospital research ethics committee.

## Results

### Development of hepatic liver organoids

We generated human liver organoid cultures from normal liver following liver resection (n=9) and core biopsy (n=3) samples using the method previously described by Hutch *et al* (Figure 1A). Patient and sample characteristics are provided in supplementary materials (Table 3). In brief, collagenised liver fragments were embedded in Matrigel and selective isolation of ductal stem cells was achieved using Isolation Medium (IM) that consisted of Expansion Medium (EM) supplemented with conditioned media containing noggin, Wnt3a and Y-27632 (see materials and methods for specific growth factors). Despite the small amount of starting material from core liver biopsies, we observed that organoids of a cystic nature grew to confluency within 2-3 weeks from isolation with 100% success rate while the success rate for organoids derived from resected liver were not as efficient at 75%. Undifferentiated organoids could be maintained for more than 6 months with weekly passaging. To determine if the organoids develop liver organ phenotype, we assessed the sequential morphological changes throughout differentiation. During the isolation and expansion phase the undifferentiated organoids assumed cystic structures of varying diameter (Figure 1B UO), however, following addition of expansion media the organoids assumed a branching “tree in buds” appearance over the course of 20 days with changes clearly visible as early as 5 days post differentiation (Figure 1B). Histological analysis (H&E staining) indicated a change from a ductal-like phenotype with single-layered epithelium to a multilayered hepatocyte-like phenotype following differentiation (Figure 1C). This differentiation process was accompanied by hepatocyte polarisation as evidenced by visualisation of Zona Occludens-1 (ZO-1: tight junction marker). There was a clear redistribution of ZO-1, in which ZO-1 is present on the complete outer membrane of undifferentiated organoids to a punctate localisation presumably at tight junctions and interconnecting structures similar to bile canaliculi in the differentiated state (Figure 1D). The ultrastructure of liver organoids examined under transmission electron microscopy showed evidence of hepatocyte phenotypes with liver microarchitecture such as hepatocyte lipid vesicles, microvilli and tight junctions bordering the bile canaliculi and glycogen granules that suggests the development of a hepatocyte phenotype (Figure 1E).

**Figure 1.**
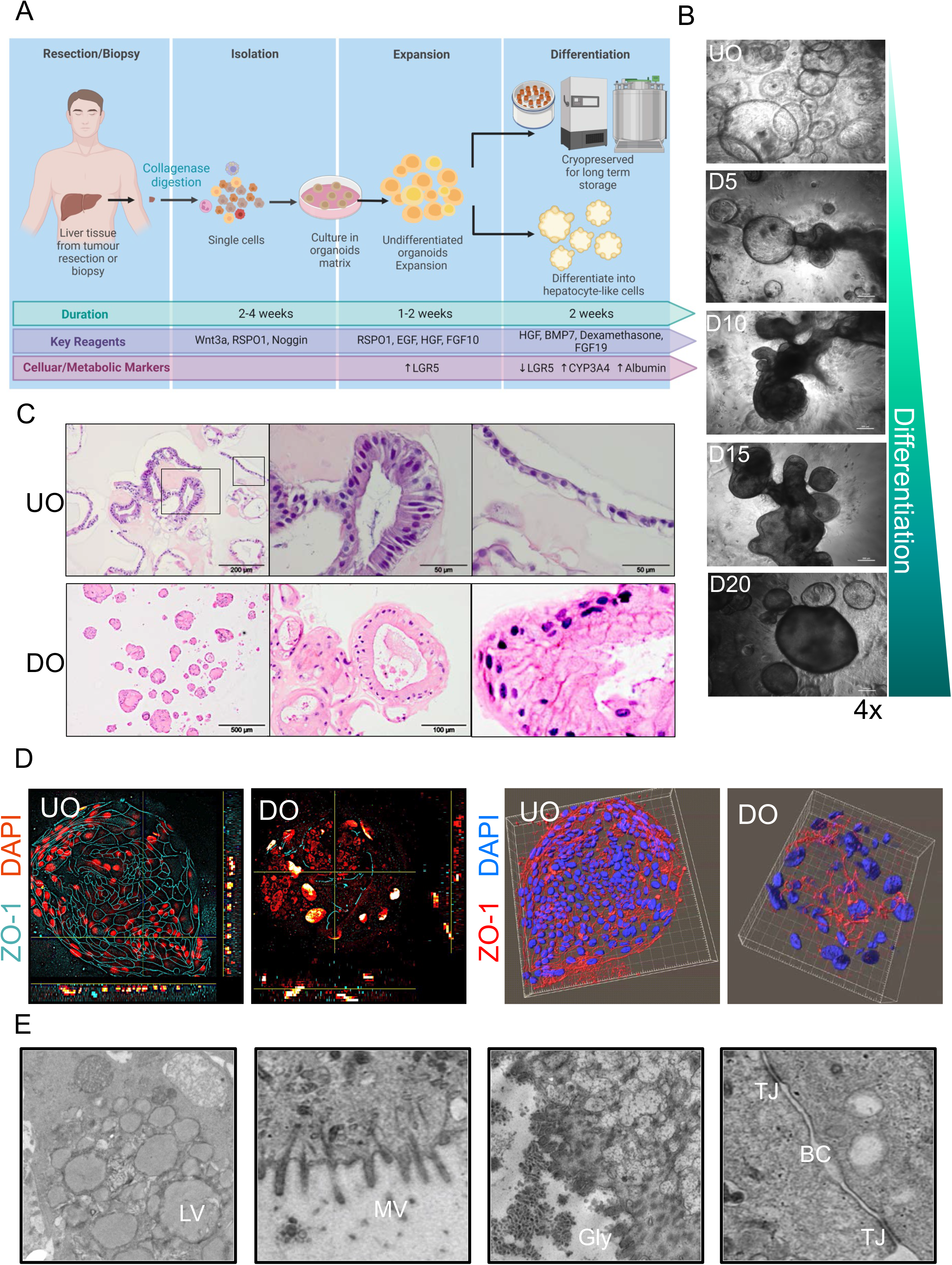
Development and characterisation of hepatic liver organoids. (A) Schematic illustration of LGR5 positive liver organoids isolation and differentiation processes (created with Biorender). (B) Brightfield microscopy of organoids from undifferentiated (UO) to differentiated organoids (DO), 4x magnification, scale bar = 200μm (C) H&E staining of UO and DO (D) Left column: 3-slide view of organoids stained with Zona-occluden-1 antibody and DAPI. Right column: 3D image deconvolution using Imaris (Bitplane). (E) Transmission Electron Microscopy (TEM) images showed cross section of UO and DO, liver microarchitectures (tight junctions [TJ] as indicated by green arrows, microvilli [MV]) and hepatocyte phenotypes (glycogen). N (nucleus), scale bar = 2μm.

**Table 3.** Patient Demographics

| Age | Gender | Disease | Recent<br>Chemotherapy | Cirrhosis | Procedure | Weight (g) |
| --- | --- | --- | --- | --- | --- | --- |
| 43 | M | Metastases | Y | N | Resection | 0.55 |
| 53 | M | Gall bladder CA | N | N | Resection | 0.97 |
| 67 | M | Metastases | N | N | Resection | 3.9 |
| 69 | M | Metastases CRC | N | N | Resection | 2.0 |
| 71 | M | Gall bladder CA | N | N | Resection | 2.7 |
| 72 | M | Metastases | N | N | Resection | 1.14 |
| 62 | M | HCC | N | Y | Resection | 2.04 |
| 57 | F | Abnormal Liver<br>Function | N | N | Biopsy | 0.02 |
| 64 | M | NAFLD | N | N | Biopsy | 0.05 |
| 26 | M | HCC | N | N | Biopsy | 0.03 |
| 62 | M | Metastases | N | N | Resection | NA |
| 67 | M | Metastases | Y | N | Resection | 2.03 |

### Liver organoids exhibit a mixed cell phenotype

Next, we examined whether differentiation associated changes to organoid structure was accompanied by gene expression changes consistent with a hepatocyte phenotype. Using qRT-PCR, metabolic markers of mature hepatocytes were assessed in undifferentiated and differentiated organoids isolated from three patients and the liver tissue from which the organoids were derived.

In comparison to undifferentiated organoids, organoids incubated in differentiation media for greater than 10 days revealed mRNA expression of the mature hepatocyte markers albumin, CYP3A4 and CYP2B6. In contrast and as expected, mRNA for the stem cell marker, LGR5 was only detected in undifferentiated organoid cultures. The hepatocyte marker (HNF4a) and ductal cell markers (KRT19) were comparably expressed at the mRNA level, before and after differentiation, suggesting retention of bi-cellular phenotypes. Using immunodetection we detected the ductal marker SOX9 and HNF4a in both undifferentiated and differentiated organoids, while albumin expression was only seen in differentiated organoids (Figure 2B). To explore the relationship between liver organoids and the tissue of origin, we performed RNASeq analysis of differentiated and undifferentiated organoids and their cognate liver tissue from 4 donors. Analysis of transcriptomic abundance revealed clustering of samples based on gene expression of liver metabolic markers with undifferentiated organoids and differentiated organoids more similar than cognate liver tissue. This is not unexpected given the significant number of hepatocytes present in the tissue samples, however it does indicate that the differentiation process drives a gene expression profile in differentiated organoids that is consistent with a hepatocyte phenotype with increased expression of Albumin, CYP3A4, CYP2C19 and a decrease in LGR5 mRNA expression. Collectively these results indicate that we have generated liver organoid cultures that consist of a mixed cell population representing mature hepatocytes and ductal cell phenotypes. Interestingly, distinct donor variation in gene expression was seen among the differentiated organoids despite similar culture and differentiation conditions (Figure 2C).

**Figure 2.**
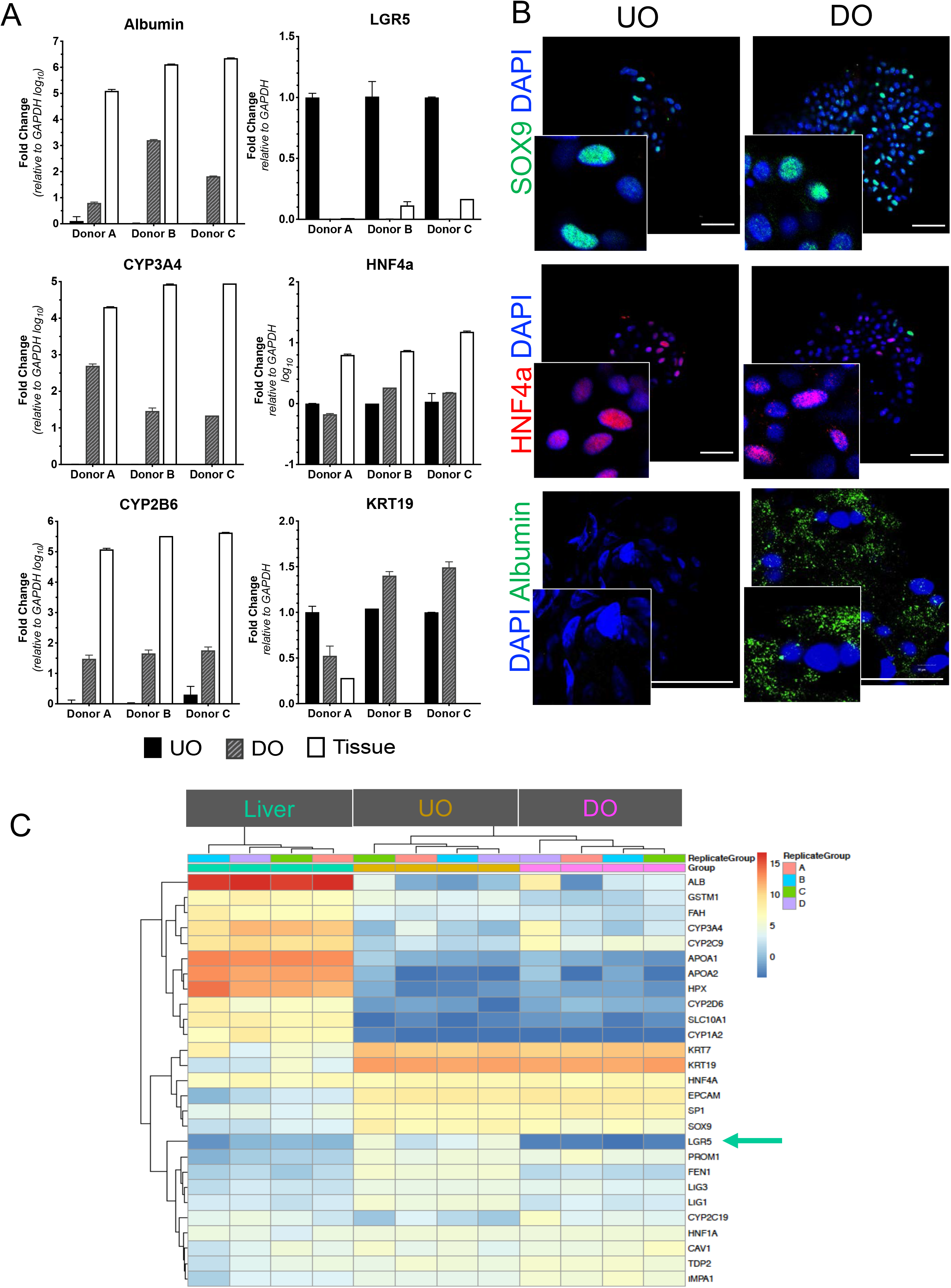
Liver organoids exhibit Interpatient variation in metabolism. (A) qRT-PCR of metabolic and cellular markers for three corresponding sets of undifferentiated organoids (UO), differentiated organoids (DO) and liver tissue (Fold change, relative to GAPDH, Mean±SEM). (B) Representative immunofluorescence staining of UO and DO for ductal marker (SOX9), hepatocyte marker (HNF4a) and albumin (20x magnification). Zoom images are shown as inset. (C) Heat map comparison of metabolic and cell signature transcriptomes from RNA-seq of 4 UO, DO cultured under similar conditions with corresponding liver tissue. (D) RNA sequencing analysis of host factors involved in HBV pathways (KEGG) for UO, DO and corresponding parental liver tissue from 4 donors.

### Liver organoids express the HBV entry receptor NTCP

To assist in downstream HBV infection studies, we next examined the temporal and spatial relationship of the HBV entry receptor expression, NTCP, throughout differentiation. The temporal development of differentiation was monitored through detection of Albumin mRNA that revealed a steady and sustained increase at day 10 post differentiation process (Fig 3A). NTCP mRNA expression while low in undifferentiated organoids did not appreciably increase until day 15, consistent with hepatocyte differentiation (Fig 3A&B). Interestingly, expression of NTCP mRNA was also donor dependent and warrants further investigation to determine if this observation correlates to *in vivo* expression and HBV infection. Next, we assessed if NTCP mRNA expression translated to functional receptor expression at the hepatocyte surface. In addition to liver organoids assuming a polarised phenotype following differentiation, inverted polarisation phenotype has been described in other types of organoids, posing potential challenges to HBV access to infection.^23^ Confocal microscopy revealed that NTCP expression was detected at the cell surface and between adjoining cells, distant from ZO-1 that is localized to apical surface (Figure 1D). Using a fluorescent labelled HBV PreS1 peptide that detects functional NTCP expression, we identified that NTCP is not expressed at the cell surface until day 10-15 following differentiation, consistent with increased mRNA expression discussed above. Interestingly, there is a low level of NTCP expression prior to day 10 although it remains intracellular suggesting that NTCP expression at the cell surface is dependent on the differentiation status of the hepatocyte. NTCP expression localised to the organoid surface most likely reflects a limit of penetration of the HBV PreS1 peptide. These results clearly reveal functional expression of NTCP on the hepatocyte surface in differentiated human liver organoids.

**Figure 3.**
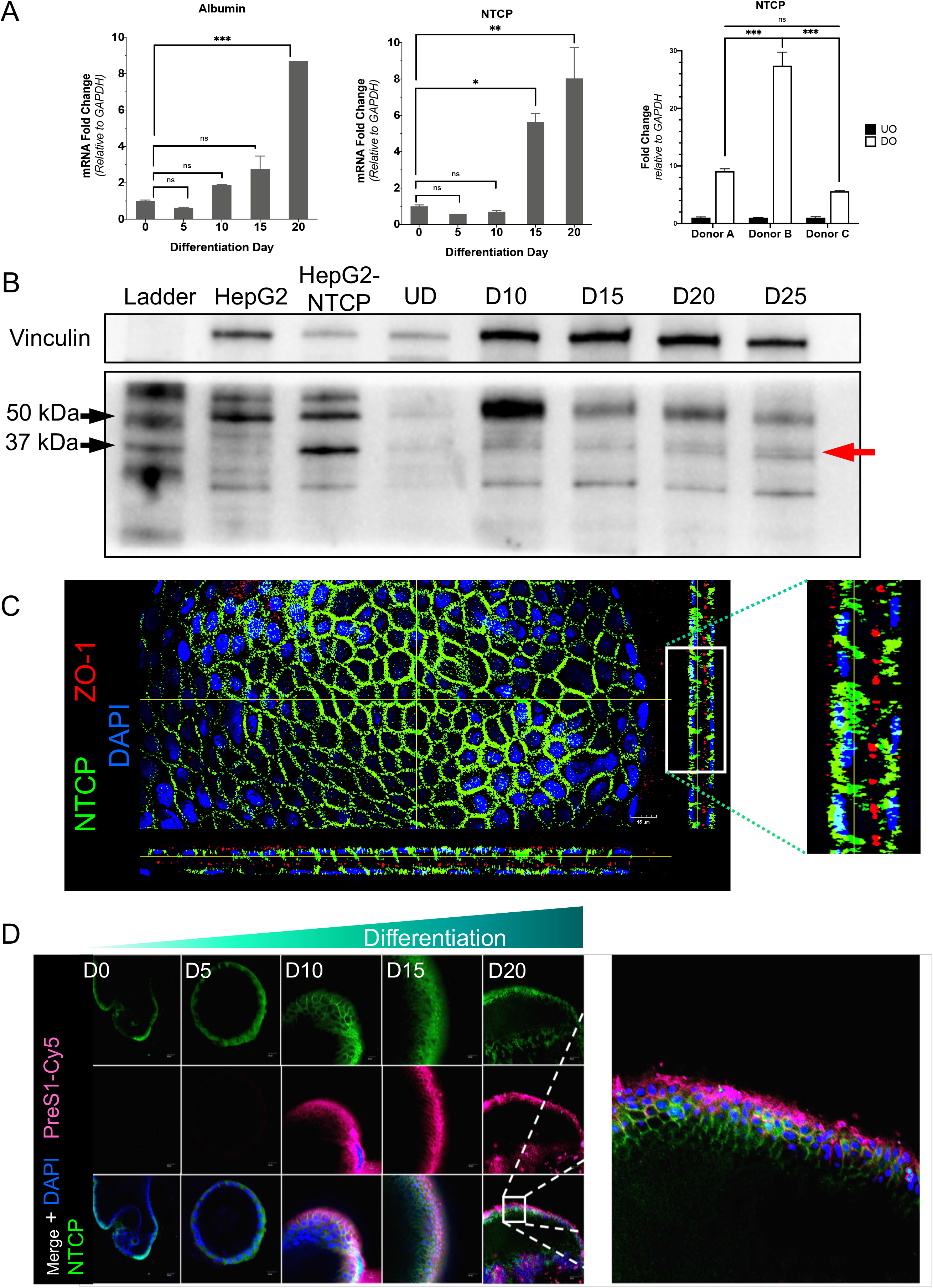
Liver organoids express functional HBV entry receptor NTCP upon differentiation. (A) Quantification of albumin and NTCP mRNA with qRT-PCR throughout differentiation (Mean±SEM, *P<005; ** P<0.01, ***P<0.001). (B) Western blot analysis of NTCP expression at indicated time point of differentiation, vinculin as expression control. (B) qRT-PCR of three corresponding UO and DO at differentiation day 15 (Fold change, relative to GAPDH, Mean±SEM, 2-way ANOVA). (C) Immunofluorescence staining of DO at day 15, cross sectional views show distribution of NTCP (green), ZO1 (red) and DAPI (blue). Scale bar = 15μm. (D) (E) Immunofluorescence staining of organoids with NTCP antibody and fluorescent tagged PreS1 throughout differentiation at 20x magnification.

### Human liver organoids support HBV infection and replication

Having established that liver organoids express functional NTCP at the hepatocyte surface we next investigated if organoids were permissive to HBV infection. Liver organoids from donors were grown in DM for 15 days prior to infection with recombinant HBV derived from HepAD38, a cell line stably expressing HBV or HBV patient positive plasma. Initial experiments using plasma derived HBV (genotype B, 1000 genome equivalents) was used to infect 15 day differentiated organoids, however, while we could detect HBcAg and HBsAg positive cells, these were scant using multiple different patient organoids (results not shown). To increase the efficiency of infection we adopted a method in which organoids were removed from the Matrigel, washed and incubated with plasma or cell culture derived HBV and subjected spinoculation at 300xg for 1hr. HBV replication was initially validated by visualisation of HBcAg expression by Immunofluorescence confocal microscopy (Fig 4A). Immunostaining following infection with cell culture derived HBV revealed staining for HBcAg 4 days post-infection with an inoculum of at least 200 genome copies/cell. As expected HBcAg expression was cytoplasmic and was present in distinct foci across the organoids that is similar to the foci of infection observed in primary hepatocytes (Fig 4B).^24^ Interestingly, the use of HBV positive plasma as inoculum resulted in infection with as low as 50 GEq/ml. We also validated and quantified HBV replication through investigation of intracellular HBV pre-genomic RNA, HBV pre-genomic and subgenomic RNA (total RNA), covalently closed circular DNA (cccDNA) and HBV DNA and HBeAg and HBsAg in the supernatant over a 2-week period (Fig 5). In all cases heat-inactivated (HI) HBV acted as a control for input inoculum. In all investigations we noted a significant increase in pgRNA, total RNA, cccDNA, extracellular HBV DNA and HBeAg/HBsAg across 3 organoid donors in comparison to the HI control. These results indicate that differentiated liver organoids are capable of supporting HBV infection and replication when challenged with either cell culture or patient derived HBV. The detection of extracellular HBV DNA and HBsAg suggests release of infectious virions, however, future studies will be required to determine if supernatant from infected organoids can infect naïve organoid cultures. Interestingly differences in donor efficiency of infection were observed that is consistent with variation in HBV permissiveness in primary human hepatocytes and most likely in natural infection ^25^. Collectively these results provide evidence that human liver derived organoids provide a platform to study not only HBV replication in a primary cell setting but also the identification of host factors and the role that genetic variation plays in infection outcome.

**Figure 4.**
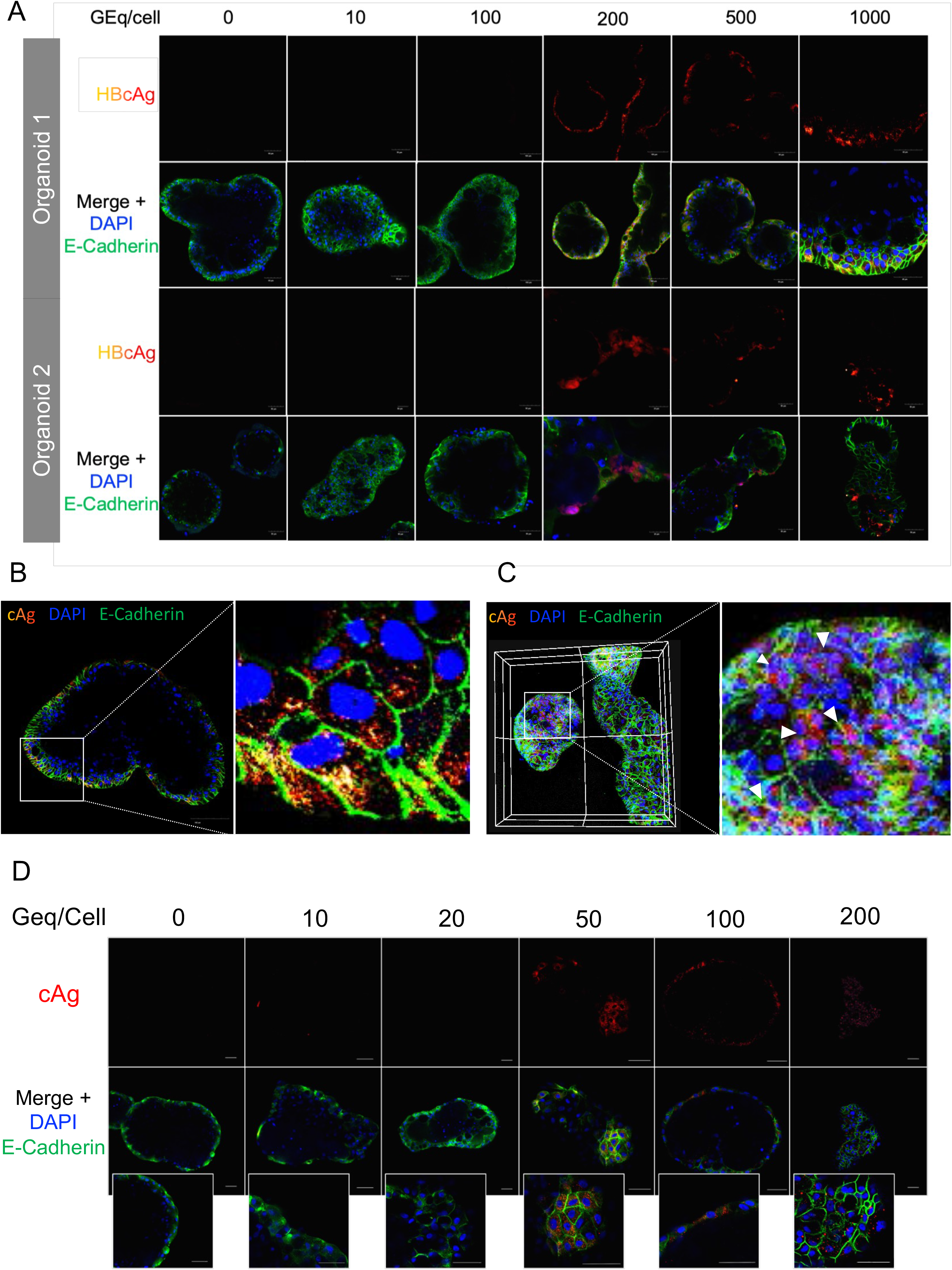
Differentiated human liver organoids are permissive to HBV infection. (A) Representative confocal microscopy images of two DO infected with cell culture-derived HBV using spinoculation at different GEq/cell (20x magnification) at 4dpi. (B) Immunofluorescence staining for cell-adhesion protein, E-Cadherin (green) and HBcAg (red/yellow) showed cytoplasmic staining of infected organoids (20x magnification). (C) Three-dimensional view of confocal images for organoids infected with HBV. White arrows indicate infected foci (zoom image). (D) Confocal microscopy images of organoids stained with HBcAg 5days post-infection with plasma HBV (genotype C) at different inoculums. Scale bar =20μm.

**Figure 5.**
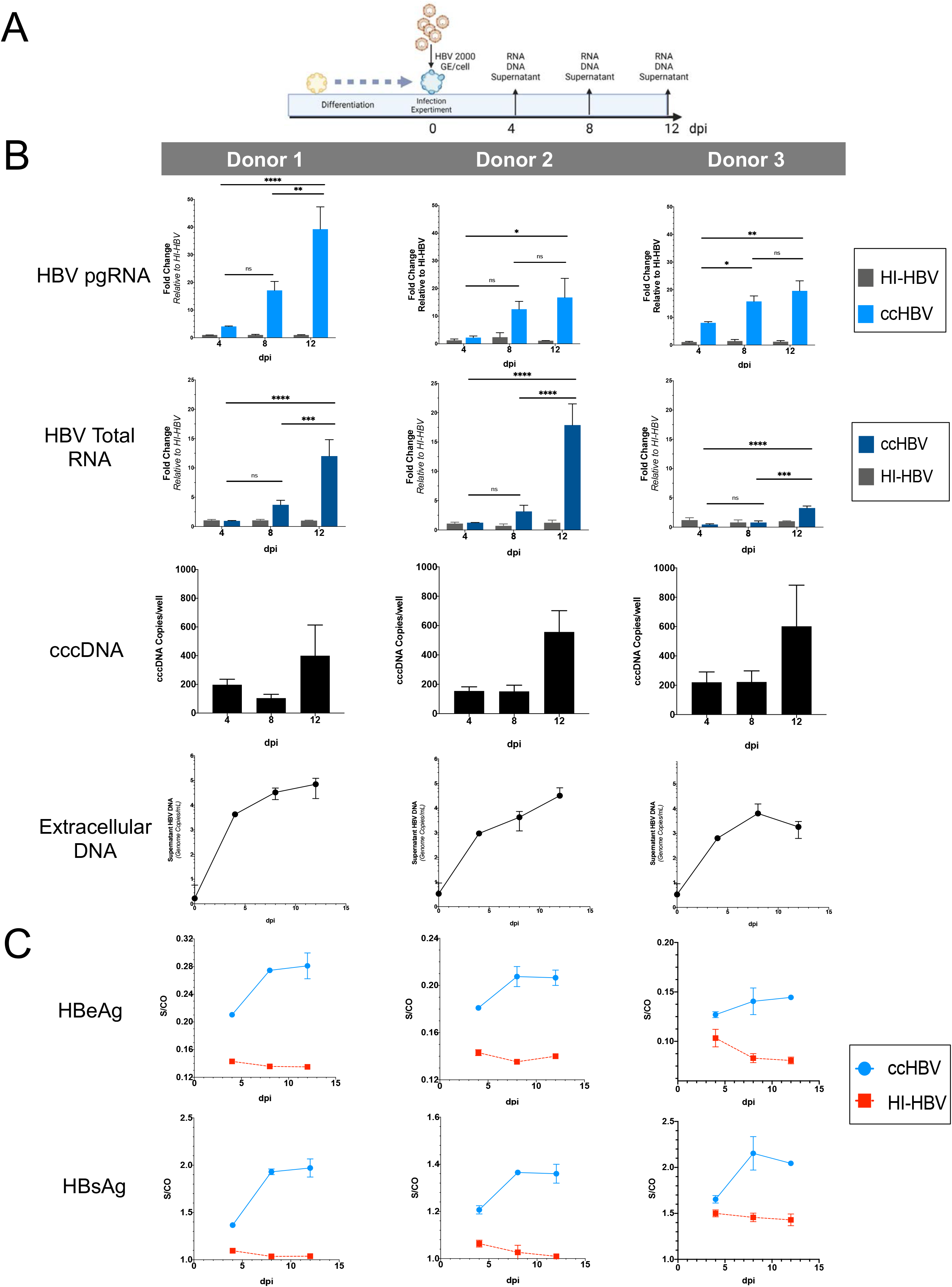
Differentiated human liver organoids support HBV replication. (A) Schematic representation of HBV infection experimental procedure and timeline for organoids derived from three separate donors. (B) HBV replication in DO was assessed with qPCR at indicated timepoints for HBV pgRNA, total RNA, cccDNA and extracellular HBV DNA (*P<005; ** P<0.01, ***P<0.001, 2-way ANOVA Mean±SEM) (ccHBV= cell culture derived HBV and HI-HBV = heat-inactivated HBV). (C) Temporal HBeAg and HBsAg excretions in supernatant were determined using ECLIA (S/CO, Mean±SEM).

### Application of Human Liver organoids in assessing antiviral drug responses

Having established that human liver organoids are permissive to HBV infection we next examined if they were suitable for anti-viral drug testing. Myrcludex B is an HBV entry inhibitor that blocks viral entry through interaction with the HBV receptor NTCP and has recently been licenced as a therapeutic for HBV and HDV.^26^ Following infection of organoids with patient derived HBV genotype B and treatment with Myrcludex B either 1 day pre- or 1 day post-infection, antiviral activity was assessed by detection of HBV total RNA and confocal immunofluorescence microscopy to detect HBcAg at 21 days post infection (Fig 6A). Consistent with the mechanism of action of Myrcludex B as an entry inhibitor, pre-infection Myrcludex treatment resulted in significant reduction of HBV RNA and HBcAg whereas post-infection treatment with Myrcludex B did not as replication had been established (Fig 6A).

**Figure 6.**
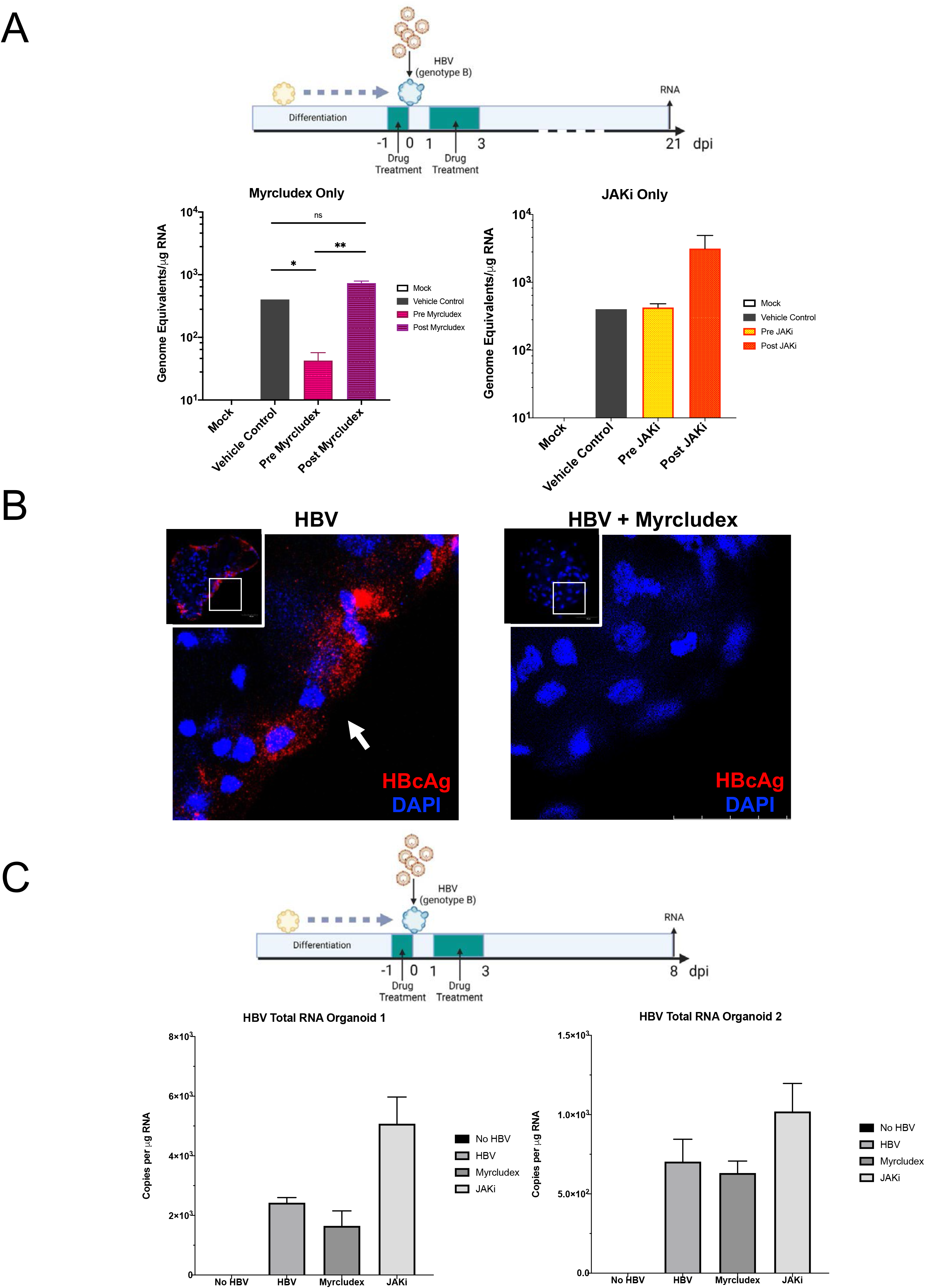
Human liver organoids shows drug responsiveness to entry inhibitor and JAK1/2 inhibitor. Drug treatment of liver organoids infected with plasma HBV (genotype B) for 21 days. (A) Schematic timeline of experiment and qRT-PCR of infected organoids, normalised to total RNA, (*P<005; ** P<0.01, 2-way ANOVA, Mean±SEM) for Myrcludex B and JAKi (Baricitinib). (B) Corresponding immunofluorescence for HBcAg in the same experiment. (C) Schematics of treatment timeline for two organoids in a shorter time frame and qRT-PCR of infected organoids, normalised to total RNA, expressed as Mean±SEM.

Baricitinib is a potent JAK1/2 inhibitor used to dampen the interferon response and is used clinically in rheumatoid arthritis (RA). Baricitinib use in patients with RA and HBV infection in many cases results in increases in HBV replication and suggests that HBV replication is sensitive to interferon dependent or independent activation of Jak1/2 and the innate antiviral response. Interestingly, post-infection treatment of 2 donor organoids with the JAK inhibitor resulted in an increase in HBV RNA that was not observed with pre-infection treatment (Fig 6A & 6C). This suggests that HBV replication is sensitive to innate activation of antiviral responses and is consistent with studies showing that HBV can be cleared from the liver in a cytokine-mediated non-toxic manner ^27, 28^ and as a result of interferon dependent antiviral responses.^29^

### Human liver organoids demonstrate a robust ISG response to interferon-α and dsRNA mimics but not HBV infection

The innate immune response to viral infection is the first line of defence against invading pathogens and is crucial for the outcome of viral infection and shaping a robust adaptive response. Previous studies have suggested that HBV acts as a stealth virus that is undetected by cellular pattern recognition receptors resulting in a blunted antiviral innate response.^30^ Our HBV organoid infection model thus provides a suitable system to evaluate HBV induced innate responses in physiologically relevant cells. However, before we investigated this response, we investigated the innate immune competency of our human liver organoids. Organoids from a number of different donors, either undifferentiated or differentiated were stimulated with interferon-α or PolyI:C and ISG mRNA expression evaluated. Timepoint experiments revealed comparable upregulation of mRNA for the ISGs IFITM1, ISG15 and viperin following stimulation with 1000U/mL of interferon-α for both undifferentiated and differentiated organoids (Figure 7A). Even at lower concentrations of interferon-α (10U/mL and 100U/mL), we noted significant upregulation of mRNA for ISGs and the proinflammatory cytokine, CXCL10 (Figure 7B). While only a small number of donors were investigated it was immediately apparent that the response to interferon-α was donor dependant (Fig 7C). For example, mRNA expression of the ISG IFITM1 in donor 3 was significantly reduced compared to donors 1 and 2 while the converse was true for CXCL10. Similar to interferon-α, we also noted a 2-4 log induction of ISG mRNA following stimulation of undifferentiated and differentiated organoids with the RNA viral genome mimic PolyI:C (2μg/mL) that activates the innate response through cytosolic (RLRs) and endosomal TLRs. Collectively these results suggest that human derived liver organoids retain innate immune factors for detection and response to viral infection. Next, we assessed the innate response to HBV infected organoid cultures. Three individual donor human liver organoids, the same that were characterised for HBV replication in figure 5, were infected with HBV and monitored for innate immune activation over a course of 12 days. Heat inactivated HBV was used as a control. Consistent with previous reports we noted no induction of interferon-β production or ISG expression across all donors even in the face of productive HBV infection confirming that, at least in human derived liver organoids, HBV does not activate innate immune responses.

**Figure 7.**
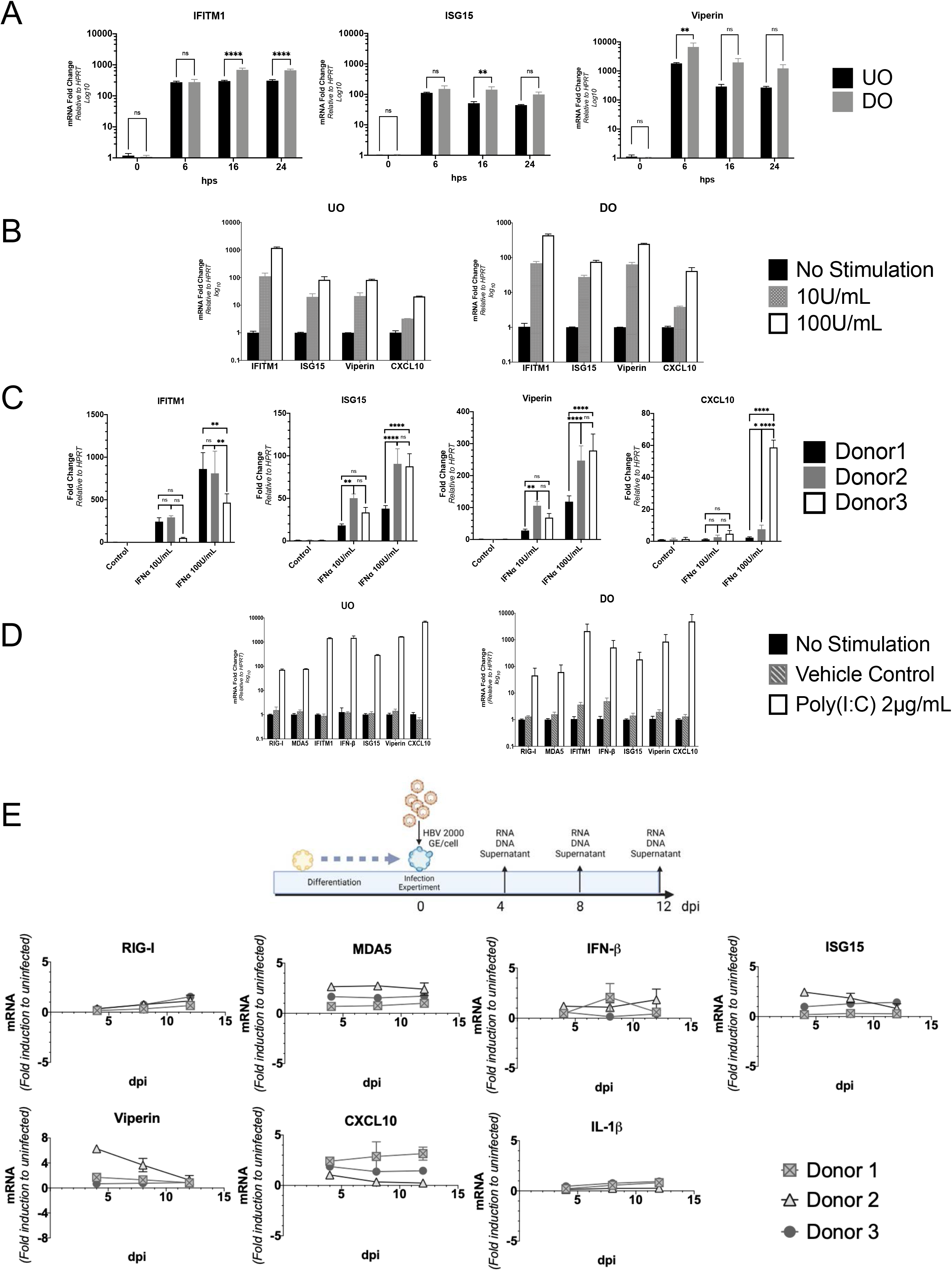
Human liver organoids demonstrate a robust ISG response to interferon-⍰ and RNA mimics but not HBV infection. (A) Temporal fold induction of ISGs in UO and DO at 6, 16 and 24h post-stimulation with interferon-α at 1000U/mL (n=2). (B) Fold induction of ISGs at 24h post-stimulation with two concentrations of interferon-α. (C) Fold induction of ISGs in DO derived from three separate donors at 24h post-stimulation with interferon-α. (D) Fold induction of viral sensing and anti-viral effector ISGs in UO an DO 24h post-stimulation with PolyI:C at 2μg/mL and (E) following HBV infection among three organoids at 1000 GEq/cell. mRNA expression relative to HPRT and normalised to uninfected controls at each specific time points. (*P<005; ** P<0.01, 2-way ANOVA, Mean±SEM)

## Discussion

International efforts are currently underway to achieve either complete or functional cure for HBV infection. This will ultimately be only achieved through the development of novel therapeutic agents used in new treatment regimens or in combination with current therapeutic strategies. Emerging therapeutics in preclinical and Phase I/II clinical studies range from direct-acting agents that target the HBV life cycle (e.g. anti-HBV RNA, entry inhibitors, RNA translation inhibitors, capsid and HBsAg inhibitors)^31–33^, enhancement of the innate immune pathways (RIG-I NOD2, TLR7 & TLR8 agonists)^27, 34^, modulation of host-factors (apoptosis inducer, ciclofillin inhibitor)^35, 36^ and gene editing technology (CRISPR/Cas9, ARCUS platform)^37^. However, assessment of the efficacy of these emerging novel therapeutic strategies and their impact on the hepatocyte will require an *in vitro* model system that not only supports the full lifecycle of HBV but also accounts for the hepatocyte phenotype and individual genetic patient differences. Primary human hepatocytes may fulfil some of these roles, but rapid dedifferentiation as a result of major transcriptomic alterations of genes involved in TCA cycle, mitochondrial dysfunction and oxidative phosphorylation can occur as early as 4 hours after isolation from liver ^38, 39^ while their limited proliferative capacity, availability and cost restricts their use for viral infection and cell response studies.

To overcome the current limitations of *in vitro* HBV model systems we developed LGR5+ stem cell derived human liver organoids originally described by *Hutch et al* that allow long-term expansion in the undifferentiated form with genetic stability^5^. The process of organoid generation from liver derived stem cells requires stimulation of the Wnt-β-catenin pathway, which recapitulates the regenerative process of liver lobules from stem cells that reside in close proximity to the central veins, following liver injury.^40^ Indeed, following differentiation *in vitro*, they express both mature hepatocyte and ductal markers and transfer of organoids in a mouse transplantation model, resulted in successful engraftment and development of mature hepatocytes with suppressed ductal phenotypes.^41^ Using this physiologically relevant model system, we successfully isolated and propagated human liver derived organoid cultures from both core biopsy and resected liver tissue with a long-term view to examine HBV infection dynamics in different donors and assess relevant cellular responses to HBV infection.

Having established that our liver organoid cultures displayed a mixed cell phenotype of hepatocytes (i.e., ALB+ve, CYP3A4+ve, CYP2C9+ve and LGR5-ve) and to a lesser extent bile duct epithelial cells (KRT19 +ve), consistent with previous work form Hutch *et al*, we next evaluated their ability to support HBV replication. Organoids derived from “normal” liver tissue from 3 different donors were infected with recombinant HBV derived from HepAD38 cells or HBV derived from patient plasma (genotype B) and over a 12-day time course revealed expression of HBV pre-genomic RNA, total RNA and ccDNA and extracellular HBV DNA and expression of HBeAg and HBsAg. Moreover, immunofluorescence microscopy revealed cytoplasmic HBV core antigen expression in organoid hepatocytes. These results collectively indicate that human liver organoids support the complete HBV life cycle and are a viable platform for the study of the molecular mechanisms of HBV replication and the cellular response in a primary cell setting.

NTCP (SCL10A1) is an essential HBV entry receptor expressed at the basolateral surface of the hepatocytes^42^ and it has been suggested that primary human hepatocytes lose their permissiveness to HBV infection as a result of NTCP expression following isolation and differentiation in culture.^43^ NTCP is a transmembrane glycoprotein with C-terminal located at the extracellular portion and N-terminal located at the intracellular portion of hepatocytes^44^. In our studies expression of NTCP mRNA correlates with differentiation status, and while low level expression of NTCP was noted in undifferentiated organoids this was predominantly intracellular and it was not until day 10 post differentiation that NTCP was expressed at the cell surface as determined using a HBV PreS1 peptide binding assay. This suggests that hepatocyte differentiation is essential for functional NTCP expression and for productive HBV binding/entry in liver organoids. Consistent with this, infection of liver organoids was dependent on NTCP as revealed by treatment of organoids with the NTCP competitive HBV entry inhibitor Myrcludex-B (see below). Interestingly, low level intracellular expression of NTCP in undifferentiated organoids suggests possible post-translational modification of NTCP in mature hepatocytes and cell surface expression of the HBV binding domain. Importantly, in our differentiated liver organoids, NTCP is expressed on the external surface to allow HBV binding and entry. We also noted a significant difference in NTCP expression at the mRNA level between 3 donors suggesting possible individual variance in NTCP expression that may impact susceptibility to HBV infection. However, our infection data from the 3 donors suggests that this may not be the case and recent reports have suggested that higher expression of NTCP in liver organoid cultures does not result in more efficient HBV infection.^45^ Further experiments are required to explore genetic differences in HBV infection susceptibility.

Given that adult stem cell derived human liver organoid cultures can be readily isolated from individual patient livers and are amenable to biobanking suggests that they are a suitable platform for the study of existing and novel inhibitors of HBV replication that may inform genetic susceptibility to HBV antivirals and primary hepatocyte cellular toxicity. To this end we further assessed the antiviral activity of Myrcludex-B, a licenced HBV entry inhibitor that competitively binds the cellular HBV receptor NTCP, either in a pre or post infection scenario. As expected, treatment of liver organoids prior to HBV infection resulted in a significant decrease in HBV RNA and complete loss of expression of HBV core antigen in infected hepatocytes, while addition of Myrcludex-B following infection had no impact on HBV infection consistent with its viral entry mode of action. Baricitinib is an immunosuppressive oral selective Jak1 and Jak2 inhibitor and reactivation of HBV replication is a well-recognised complication of patients receiving Baricitinib, especially those with rheumatoid arthritis.^46^ Baricitinib treatment of HBV infected organoids resulted in an increase (approximately 1 log) in HBV RNA suggesting suppression of the tyrosine kinases Jak1 and Jak2 can increase HBV genome replication. The mechanism of this HBV increase is unclear in our organoid model, however given that the JAK-STAT signalling pathway plays a role in signal transduction for several key cytokines, a number of mechanisms are possible. Interleukin-6 plays an important role in hepatocyte homeostasis and its impairment can lead to reduced innate responses to viral infection. Conversely, disruption of interferon signalling, which is important for an antiviral state through production of antiviral ISGs, may explain the increase in HBV replication following inhibition of Jak1 and 2 by Baricitinib. Collectively, our human liver organoid HBV infection model now provides a model system to dissect the molecular mechanisms responsible for a new wave of HBV antivirals.

Hepatitis A and C virus induce strong innate antiviral immune responses in hepatocytes following infection and in response, these viruses have evolved elaborate mechanisms to combat the antiviral response. However, the ability of HBV to induce a similar robust antiviral response is controversial with inconsistent reports on the ability of HBV to activate or evade the innate antiviral response.^30, 47^ This is primarily due to the limitations of model systems used to investigate HBV replication such as hepatoma derived cell lines (i.e., HepG2) that have defects in innate immune sensing and associated signal transduction pathways. However, there is evidence that HBV can activate an antiviral innate response in physiologically relevant culture systems. Micro patterned cocultures in which primary human hepatocytes are cocultured with stromal fibroblasts are permissive for HBV infection for up to 3 weeks and pluripotent stem-cell derived hepatocytes (iHeps) induce an interferon and antiviral ISG response. Furthermore, interferon can degrade cccDNA from the HBV infected liver and in a small subset of individuals, pegylated interferon can eliminate HBV, while in HBV infected chimpanzee’s HBV replication is controlled by a type I interferon response. ^27, 48^ However, in contrast, HBV infection of PHH neither activates of inhibits pattern recognition receptors and thus evades type-I antiviral responses and as such has been described as a stealth virus.^49^ Consistent with the report from Lucifora *et al* we also report that HBV does not induce an appreciable innate response in patient derived human liver organoid cultures as assessed by ISG mRNA expression in response to HBV infection. It is possible that this observation may be due to an inherent inability of organoid cells to respond at the innate immune level, however stimulation of either undifferentiated or differentiated liver organoids with PolyI:C or interferon-α resulted in significant induction of the ISGs, IFITM1, ISG15, Viperin and CXCL10, similar to that of PHH, confirming their ability to mount an innate immune response. Thus, we conclude that infection of adult stem cell derived liver organoids with HBV does not induce an appreciable innate immune response consistent with HBV being a stealth virus.

While the use of hepatoma cells ectopically expressing NTCP have been useful for the study of the HBV lifecycle, they do not mimic primary hepatocytes making host viral interaction studies difficult. As such in recent years there has been a significant effort to develop human liver organoid cultures as a HBV infection model system. Induced pluripotent stem cell (iPSC) derived hepatocyte-like cells have been shown to support HBV infection, however they have limitations in that during the reprogramming process there is a high rate of chromosomal mutation that may result in aberrant phenotypes and rigorous assessment of terminally differentiated cells is required to ensure authenticity. Moreover, they are one dimensional in their genetic composition that is not representative of the general population. In contrast, adult human liver stem cell derived organoids, as described in this study, do not suffer from chromosomal aberrations and most importantly, they preserve the genetic make-up of the liver tissue from which they were derived. Furthermore, they are amendable for biobanking and long-term HBV patient derived infection studies that will ultimately help define an individual’s response to HBV infection and pave the way for a personalised medicine approach to HBV management and therapeutic strategies.

## Acknowledgements

The authors are grateful for the assistance from the hepatobiliary surgeons (Dr Paul Dolan, Dr CP Tan, Dr Mark Brooke-Smith, Dr John Chen, Dr Eu Ling Neo), theatre nurses and Dr John Bates (gastroenterologist) from the Royal Adelaide Hospital for acquiring liver tissues. The authors also thank Maria Collis from SA Pathology for coordinating tissue sectioning and Helen Beard from SAHMRI for her assistance with histology; Dr Stuart McKessar, Ms Tara Green, Mr Craig Riddle and Dr Geoffrey Higgins for their technical assistance with HBV serology. Special thanks to Ms Catherine Ferguson, A/Prof David Shaw and Dr Morgyn Warner for their assistance with HBV blood collection.

